# Bridging the dynamics and organization of chromatin domains by mathematical modeling

**DOI:** 10.1101/151563

**Authors:** Soya Shinkai, Tadasu Nozaki, Kazuhiro Maeshima, Yuichi Togashi

## Abstract

The genome is three-dimensionally organized in the cell, and the mammalian genome DNA is partitioned into submegabase-sized chromatin domains. Genome functions are regulated within and across the domains according to their organization, whereas the chromatin itself is highly dynamic. However, the details of such dynamic organization of chromatin domains in living cells remain unclear. To unify chromatin dynamics and organization, we recently demonstrated that structural information of chromatin domains in living human cells can be extracted from analyses of the subdiffusive nucleosome movement using mathematical modeling. Our mathematical analysis suggested that as the chromatin domain becomes smaller and more compact, nucleosome movement becomes increasingly restricted. Here, we show the implication of these results for bridging the gap between chromatin dynamics and organization, and provide physical insight into chromatin domains as efficient units to conduct genome functions in the thermal noisy environment of the cell.

**Extra View to:** Shinkai S, Nozaki T, Maeshima K, Togashi Y. Dynamic Nucleosome Movement Provides Structural Information of Topological Chromatin Domains in Living Human Cells. PLoS Comput Biol. 2016;12(10):e1005136. doi: 10.1371/journal.pcbi.1005136. PubMed PMID: 27764097; PubMed Central PMCID: PMCPMC5072619.

**Funding:** This work was supported by Platform Project for Supporting in Drug Discovery and Life Science Research (Platform for Dynamic Approaches to Living System) from MEXT and AMED; KAKENHI under Grant JP16H01408, JP23115007, JP23115005, JP16H04746; CREST grant (JPMJCR15G2) from JST; and Research Fellowship for Young Scientists under Grant JP13J04821, JP16J07205.

**Disclosure:** No potential conflicts of interest were disclosed.

**Acknowledgements:** We would like to thank Dr. Takashi Toda for comments regarding the manuscript.

## Introduction

The genome is not only one-dimensional information of DNA sequences, but is further three-dimensionally organized into chromatin in the cell, with effective regulation of various genome functions. Recent developments in microscopy imaging and genomic analysis have revealed detailed information on the spatial chromatin organization. With respect to local chromatin organization, it has been suggested that the nucleosomes, consisting of DNA wrapped around core histones, are irregularly folded without the regular chromatin fibers (1-5). Electron and super-resolution fluorescence microscopy studies have revealed the existence of even larger chromatin organizations, including chromatin domains (6) or chromonema fibers (7). Recently, chromosome conformation capture (3C) derivative methods have demonstrated that the mammalian genome DNA is partitioned into submegabase-sized chromatin domains, including topologically associating domains (TADs) (8, 9), and contact and loop domains (10). In addition, computational modeling methods have been developed to reconstruct the three-dimensional (3D) architecture of the genome (11, 12).

Meanwhile, chromatin itself is highly dynamic in living cells (13-20). In particular, single-nucleosome imaging in living mammalian cells has revealed random nucleosome fluctuations driven by the thermal random force (21-24). However, it is still unclear how the chromatin domains are dynamically organized in living cells whose environment is thermally noisy. Here, we highlight the implication of our study aiming to bridge the gap between the dynamics and organization of chromatin domains in living cells through mathematical modeling (24), and provide physical insight into a chromatin domain as a regulatory and structural unit.

### Physical information extracted from molecular movement

Molecules are dynamic and diffusive in living cells. Microscopic imaging technologies such as fluorescence recovery after photobleaching (FRAP), fluorescence correlation spectroscopy (FCS), and single-particle tracking (SPT) have facilitated measurements and visualization of the dynamic properties of molecules in living cells, e.g. (25). In these methods, the diffusion coefficient *D* is quantified as a measure of the mobility of the molecules. The physical dimension of *D* corresponds to (length)^2^/(time), representing the area of a molecule diffusing per unit time. In the SPT method, under the condition of *normal* diffusion, the mean-squared displacement (MSD) is proportional to both D and time t (Fig. 1). In addition, the Stokes-Einstein equation for Brownian motion, *D* = *k*_B_*T*/6*πηr*, tells us that the origin of the diffusion is inherently related to other physical parameters: temperature *T*, viscosity η of the environment, and the hydrodynamic radius r of the molecules (26). Here, *k*_B_ represents the Boltzmann constant. Therefore, by quantifying the diffusion coefficient *D*, we can obtain the other relevant physical information such as *η* and *r* via the relation above. Note that, to determine *η* or *r*, we need an estimation of either parameter beforehand; for example, in an FCS experiment for quantum dots (QDs), hydrodynamic radii *r* of the QDs are estimated by comparison with the diffusion time of standard fluorescent beads in PBS buffer solutions; and the cytoplasmic viscosity is finally determined by use of the Stokes-Einstein equation and the measured diffusion coefficients of the QDs (27). In addition, inclusion of the term *k*_B_*T* with the energy dimension represents that the thermal noise is the driving force of diffusion showing Brownian motion.

**Figure 1.**
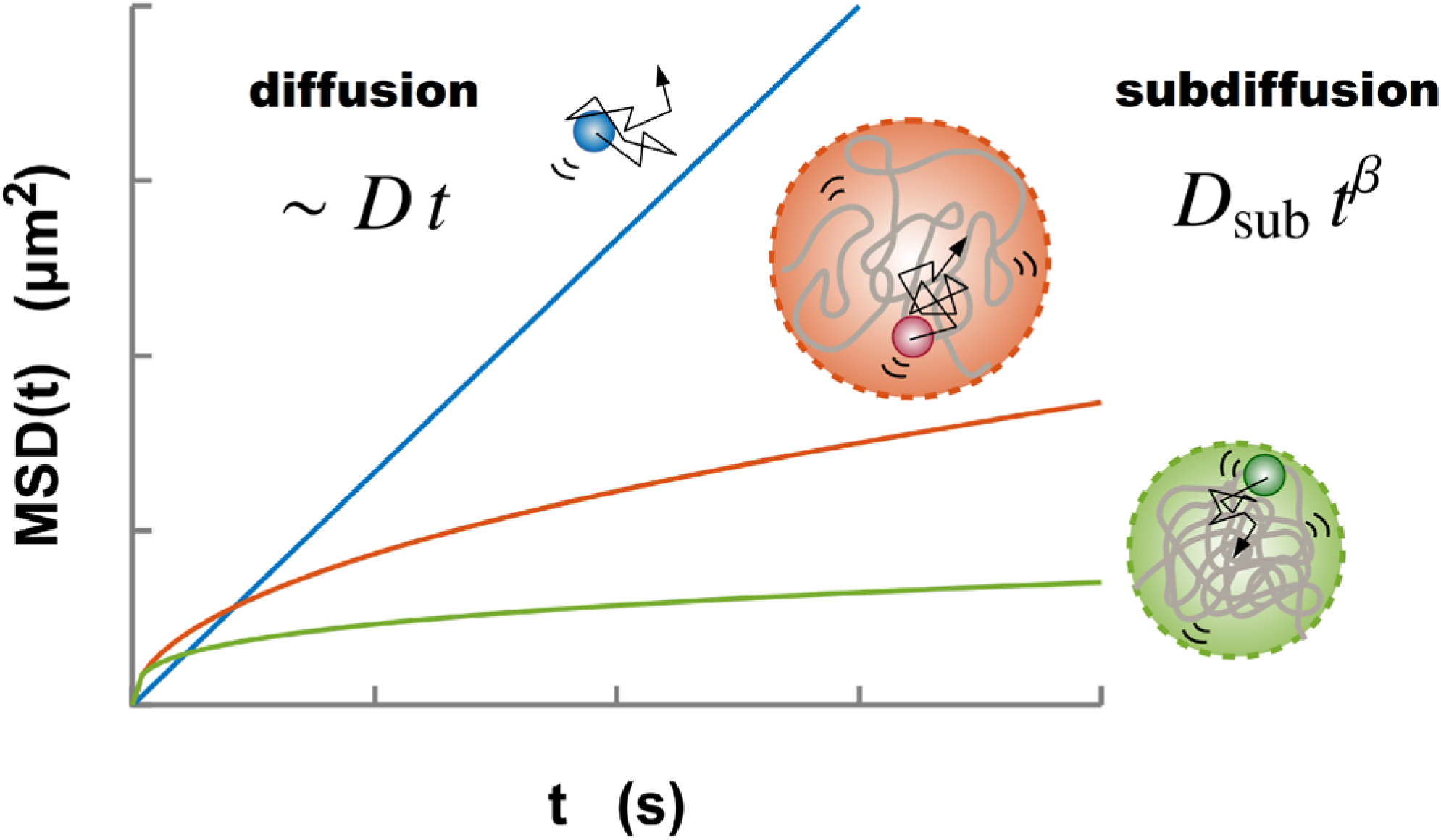
Normal diffusion (blue) of a particle driven by thermal noise characterized by the MSD, which is proportional to both the diffusion coefficient *D* and time *t*. Meanwhile, the diffusive movement of a monomer within a polymer globule represents the subdiffusion 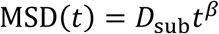 due to the structural restriction of the globule. A larger globule results in higher mobility of a monomer (orange, upper). As is shown in Eq. (2), as a polymer globule becomes smaller and more compact, the MSD also becomes smaller (green, lower).

### Can we extract physical information from chromatin dynamics?

Live-cell imaging experiments for certain chromosomal loci and nucleosomes have revealed that chromatin is highly dynamic in the interphase (13-24). To quantify these dynamics, MSD analysis is often used through SPT. Interestingly, unlike the case of normal diffusion that is proportional to time, the MSD results of chromatin show much slower diffusion with nonlinear scaling called *subdiffusion*:
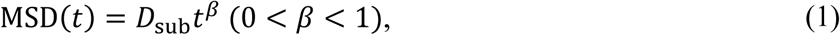

where *D*_sub_ is the coefficient of subdiffusion with the physical dimension (length)^2^/(time)*^β^*, and *β* is the scaling exponent (Fig. 1). The subdiffusive movement of chromatin has been observed generally, regardless of species and cell types (15-17, 19-24), suggesting that there must be a common principle generating the subdiffusion. The thermal noise that drives random fluctuations of chromatin in living cells is a mechanism common to the diffusion of molecules. Unlike small molecules in the nucleus, the nucleosome fiber in chromatin is a biopolymer. Therefore, the movement of the fibers can be constrained by their own organization. Accordingly, there must be a framework that unifies chromatin dynamics and their organization. As mentioned above, under a normal diffusion process, the Stokes-Einstein equation provides such a bridge for determining a physical parameter such as viscosity *η* of the environment or the hydrodynamic radius *r* of the molecules from the measured diffusion coefficient *D*. In a similar manner, mathematical modeling of chromatin dynamics should enable extraction of physical information related to chromatin organization from the dynamics.

### Mathematical model of single nucleosomes with fractal chromatin domains

In our previous work (24) to explain the subdiffusive movement of single nucleosomes, we took into account the following two assumptions: (i) each tracked single nucleosome belongs to a submegabase-sized chromatin domain in the living cell; and (ii) the effective conformational state of the chromatin domain formed by a nucleosome fiber can be characterized by using the fractal dimension *d*_f_ (Fig. 2). The nucleosome fiber was mathematically modeled as a chain of *N* monomers corresponding to nucleosomes, and the size of the chromatin domain is represented by *R*. According to the scaling law of polymers, 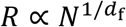, polymer conformations in 3D space are characterized by the fractal dimensions 1 ≤ *d*_f_ ≤ 3 (28) (Fig. 2). For example, *d*_f_ = 1 and 3 correspond to a straight line and the most compact state, respectively. In case where each monomer-monomer interaction is repulsive and polymer crossing is prohibited (excluded volume effect), the polymer tends to form an extended state with *d*_f_ ≅ 5/3. A polymer with *d*_f_ = 2, which is known as the ideal chain, corresponds to a random-walk conformation (Fig. 2). Although, in general, nucleosome-nucleosome interactions within a chromatin domain are thought to show complex interactions with both repulsive and attractive forces, use of the fractal dimension enables us to mathematically express the conformational state of chromatin domains without needing to identify and describe the complicated interactions.

**Figure 2.**
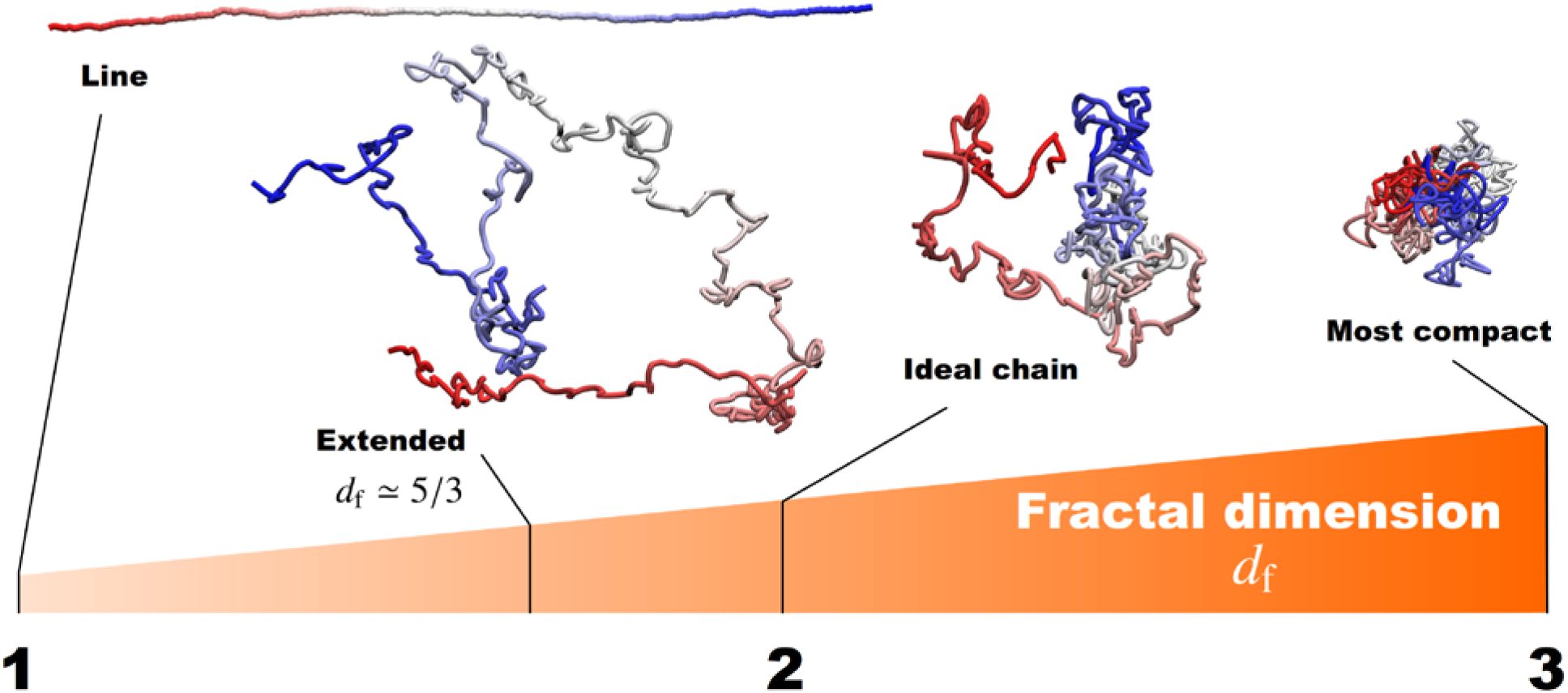
Fractal dimensions for polymers to characterize the effective conformational states. For example, *d*_f_ = 1 and 3 correspond to a straight line and the most compact state, respectively. The polymer with the excluded volume effect forms an extended state with *d*_f_ ≅ 5/3. For *d*_f_ = 2, the ideal chain is a polymer corresponding to a random-walk conformation.

Although every assumption should be verified explicitly in the future, it is nevertheless possible to mathematically describe nucleosome dynamics by adopting these assumptions (i and ii). The dynamics of each nucleosome are described by the force balance equation *F*_friction_ = *F*_fractal_ + *F*_thermal_, representing the friction force, the force required to keep a fractal domain structure formed by a nucleosome fiber, and the thermal driving force, respectively. Based on the above setting, we analytically derived the MSD in terms of polymer dynamics (24). As expected, the MSD obeys subdiffusion with the coefficient and the exponent 
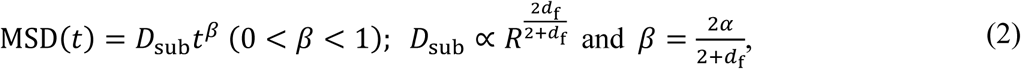

where 0 < *α* ≤ 1 is the variable related to the viscoelastic nucleus environment; we assume here that *α* is a constant value. These relations mean that the coefficient *D*_sub_ and the exponent *β* of the subdiffusive movement mutually connect to the structural parameters of the chromatin domains *R* and *d*_f_. That is, when the size R is small, the coefficient *D*_sub_ also becomes small, and thus the MSD becomes small. From another perspective, when the fractal dimension *d*_f_ increases, the exponent *β* decreases, and the MSD becomes small. Therefore, smaller and more compact chromatin domains will have smaller MSD values (Fig. 1).

### Subdiffusive single-nucleosome movement in living cells

We next applied our model to living human cells. To achieve single-particle imaging of nucleosomes in living cells, we combined an oblique illumination microscopy and labeling of histone H2B with a photoactivatable (PA)-red fluorescent protein (mCherry) (21-24, 29, 30) (Fig. 3A-C). The oblique illumination microscopy can illuminate a limited area in the nucleus with very low background noise (Fig. 3A). When we looked at the HeLa cells stably expressing H2B-PA-mCherry with the microscopy, we found that a relatively small number of H2B-PA-mCherry molecules were spontaneously and stochastically activated without an ultraviolet laser stimulation and observed as clear dots, which are suitable for the imaging (Fig. 3B). The single-step photobleaching profile of these H2B-PA-mCherry dots confirmed that each dot represents the fluorescence derived from a single nucleosome. Then, continuously fluorescent dots were tracked for a few seconds (50 ms per time frame), and Fig. 3C shows representative trajectories of the dynamic nucleosome movement in single cells. The movement in living cells was shown to exhibit random fluctuation within the range of a few hundred nanometers in a few seconds, and such restricted Brownian-like motion is thought to be caused by thermal noise (21-23). By changing the focal plane of the microscope, we could observe such single-nucleosome dynamics not only in the nuclear interior but also at the nuclear periphery (24). The MSD of the movement at each region clearly showed subdiffusion (Fig. 3D). In addition, the nucleosome movement was observed to be more mobile in the interior than at the periphery with a larger coefficient *D*_sub_ and a larger exponent *β* of subdiffusion.

**Figure 3.**
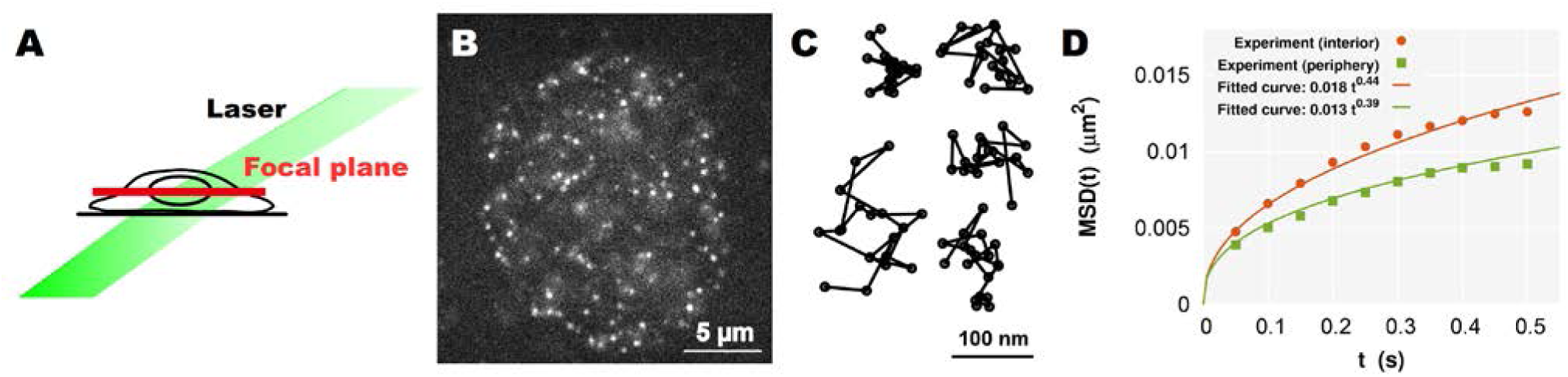
**(A)** A simple scheme for oblique illumination microscopy. An illumination laser (green) and focal plane (red) in a living cell are shown. **(B)** Single-nucleosome image of a human HeLa cell nucleus expressing H2B-PA-mCherry. Each dot represents a single nucleosome (adopted from ref. (24)). **(C)** Representative trajectories of fluorescently labeled single nucleosomes (50 ms per time frame). **(D)** Plots of the MSD at the interior (orange) and periphery (green) regions. Each plot fits well with the MSD curve for subdiffusion (Eq. (1)).

### Conversion of the mobility information of nucleosomes into structural information of chromatin domains

In Fig. 3D, the fitted curves to the MSDs at the different focal planes show the following relations
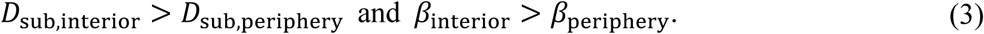

These can in turn be interpreted through our analytical result (Eq. (2)) as the structural relations
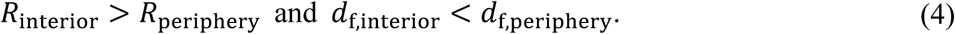

These relations are consistent with the notion that (6, 31-33) the larger and more extended chromatin domains tend to occupy the nuclear interior, whereas the smaller and more compact chromatin domains, which correspond to the heterochromatin regions, are enriched close to the nuclear periphery. Thus, we can mathematically conclude that nucleosomes in heterochromatin-rich regions show less movement, which results from the restriction of the smaller and more compact chromatin domains; although nucleosome movement will be reduced at the nuclear periphery where the heterochromatin is bound to the nuclear lamina.

More detailed quantitative estimation of these relations was carried out in ref. 24, which showed that the average size of submegabase-sized chromatin domains in living cells is in the range of 100-500 nm, and the nucleosome movement within a chromatin domain reaches thermal equilibrium in a few seconds at most. Our results suggest that dynamic nucleosome-nucleosome interactions within chromatin domains work well at a spatiotemporal scale of a few hundred nanometers and a few seconds. Thus, in terms of physics, such a chromatin domain is an efficient unit to appropriately process various genome functions in the thermal noisy environment of the cell.

### Summary and perspective

To unify chromatin dynamics and organization, we have proposed a mathematical model of a fractal chromatin domain formed by a nucleosome fiber, which was applied to the subdiffusive nucleosome movement in living cells, as measured by SPT. Through analyzing the model, we have shown that the coefficient *D*_sub_ and the exponent *β* of the subdiffusive MSD depend on the structural information of chromatin domains such as the size *R* and the fractal dimension *d*_f_. As the normal diffusion theory converts the measured diffusion coefficient into other physical parameters, these relations (Eq. (2)) can be used to provide physical insight into the subdiffusive MSD. Thus, comparison of the MSDs of nucleosome movement at different nuclear regions enables us translating the mobility information of nucleosomes into the structural information of chromatin domains: chromatin domains in the heterochromatin-rich nuclear periphery region are smaller and more compact than those in the euchromatin-rich interior region. Through more quantitative estimations, we suggest that a submegabase-sized chromatin domain of the mammalian genome is a reasonable unit to carry out various genome functions from a physics perspective.

Novel advances in specific labeling techniques, which can deal with nucleosome movement and the chromatin organization of certain specific genomic loci, will further help to uncover chromatin dynamics and their organization relating to epigenetic states and genome functions. Furthermore, based on our present model, the long-term tracking of single-nucleosome movements will allow us to ultimately unveil the larger-scale and dynamic organization of chromosomes.

## References

1. Joti Y, Hikima T, Nishino Y, Kamada F, Hihara S, Takata H, et al. Chromosomes without a 30-nm chromatin fiber. Nucleus. 2012;3(5):404-10. doi: 10.4161/nucl.21222. PubMed PMID: 22825571; PubMed Central PMCID: PMCPMC3474659.

2. Fussner E, Strauss M, Djuric U, Li R, Ahmed K, Hart M, et al. Open and closed domains in the mouse genome are configured as 10-nm chromatin fibres. EMBO Rep. 2012;13(11):992-6. doi: 10.1038/embor.2012.139. PubMed PMID: 22986547; PubMed Central PMCID: PMCPMC3492707.

3. Ricci MA, Manzo C, Garcia-Parajo MF, Lakadamyali M, Cosma MP. Chromatin fibers are formed by heterogeneous groups of nucleosomes in vivo. Cell. 2015;160(6):1145-58. doi: 10.1016/j.cell.2015.01.054. PubMed PMID: 25768910.

4. Chen C, Lim HH, Shi J, Tamura S, Maeshima K, Surana U, et al. Budding yeast chromatin is dispersed in a crowded nucleoplasm in vivo. Molecular Biology of the Cell. 2016;27(21):3357-68. doi: 10.1091/mbc.E16-07-0506.

5. Maeshima K, Rogge R, Tamura S, Joti Y, Hikima T, Szerlong H, et al. Nucleosomal arrays self-assemble into supramolecular globular structures lacking 30-nm fibers. EMBO J. 2016;35(10):1115-32. doi: 10.15252/embj.201592660. PubMed PMID: 27072995; PubMed Central PMCID: PMCPMC4868957.

6. Cremer T, Cremer M, Hubner B, Strickfaden H, Smeets D, Popken J, et al. The 4D nucleome: Evidence for a dynamic nuclear landscape based on co-aligned active and inactive nuclear compartments. FEBS Lett. 2015;589(20 Pt A):2931-43. doi: 10.1016/j.febslet.2015.05.037. PubMed PMID: 26028501.

7. Bian Q, Belmont AS. Revisiting higher-order and large-scale chromatin organization. Current Opinion in Cell Biology. 2012;24(3):359-66. doi: 10.1016/j.ceb.2012.03.003.

8. Dixon JR, Selvaraj S, Yue F, Kim A, Li Y, Shen Y, et al. Topological domains in mammalian genomes identified by analysis of chromatin interactions. Nature. 2012;485(7398):376-80. doi: 10.1038/nature11082. PubMed PMID: 22495300; PubMed Central PMCID: PMCPMC3356448.

9. Nora EP, Lajoie BR, Schulz EG, Giorgetti L, Okamoto I, Servant N, et al. Spatial partitioning of the regulatory landscape of the X-inactivation centre. Nature. 2012;485(7398):381-5. doi: 10.1038/nature11049. PubMed PMID: 22495304; PubMed Central PMCID: PMCPMC3555144.

10. Rao SS, Huntley MH, Durand NC, Stamenova EK, Bochkov ID, Robinson JT, et al. A 3D map of the human genome at kilobase resolution reveals principles of chromatin looping. Cell. 2014;159(7):1665-80. doi: 10.1016/j.cell.2014.11.021. PubMed PMID: 25497547.

11. Bau D, Sanyal A, Lajoie BR, Capriotti E, Byron M, Lawrence JB, et al. The three-dimensional folding of the alpha-globin gene domain reveals formation of chromatin globules. Nat Struct Mol Biol. 2011;18(1):107-14. doi: 10.1038/nsmb.1936. PubMed PMID: 21131981; PubMed Central PMCID: PMCPMC3056208.

12. Serra F, Di Stefano M, Spill YG, Cuartero Y, Goodstadt M, Bau D, et al. Restraint-based three-dimensional modeling of genomes and genomic domains. FEBS Lett. 2015;589(20 Pt A):2987-95. doi: 10.1016/j.febslet.2015.05.012. PubMed PMID: 25980604.

13. Heun P, Laroche T, Shimada K, Furrer P, Gasser SM. Chromosome Dynamics in the Yeast Interphase Nucleus. Science. 2001;294:2181-6.

14. Levi V, Ruan Q, Plutz M, Belmont AS, Gratton E. Chromatin dynamics in interphase cells revealed by tracking in a two-photon excitation microscope. Biophys J. 2005;89(6):4275-85. doi: 10.1529/biophysj.105.066670. PubMed PMID: 16150965; PubMed Central PMCID: PMCPMC1366992.

15. Hajjoul H, Mathon J, Ranchon H, Goiffon I, Mozziconacci J, Albert B, et al. High-throughput chromatin motion tracking in living yeast reveals the flexibility of the fiber throughout the genome. Genome Res. 2013;23(11):1829-38. doi: 10.1101/gr.157008.113. PubMed PMID: 24077391; PubMed Central PMCID: PMCPMC3814883.

16. Zidovska A, Weitz DA, Mitchison TJ. Micron-scale coherence in interphase chromatin dynamics. Proc Natl Acad Sci U S A. 2013;110(39):15555-60. doi: 10.1073/pnas.1220313110. PubMed PMID: 24019504; PubMed Central PMCID: PMCPMC3785772.

17. Lucas JS, Zhang Y, Dudko OK, Murre C. 3D trajectories adopted by coding and regulatory DNA elements: first-passage times for genomic interactions. Cell. 2014;158(2):339-52. doi: 10.1016/j.cell.2014.05.036. PubMed PMID: 24998931; PubMed Central PMCID: PMCPMC4113018.

18. Ochiai H, Sugawara T, Yamamoto T. Simultaneous live imaging of the transcription and nuclear position of specific genes. Nucleic Acids Res. 2015;43(19):e127. doi: 10.1093/nar/gkv624. PubMed PMID: 26092696; PubMed Central PMCID: PMCPMC4627063.

19. Bronshtein I, Kanter I, Kepten E, Lindner M, Berezin S, Shav-Tal Y, et al. Exploring chromatin organization mechanisms through its dynamic properties. Nucleus. 2016;7(1):27-33. doi: 10.1080/19491034.2016.1139272. PubMed PMID: 26854963; PubMed Central PMCID: PMCPMC4916879.

20. Amitai A, Seeber A, Gasser SM, Holcman D. Visualization of Chromatin Decompaction and Break Site Extrusion as Predicted by Statistical Polymer Modeling of Single-Locus Trajectories. Cell Rep. 2017;18(5):1200-14. doi: 10.1016/j.celrep.2017.01.018. PubMed PMID: 28147275.

21. Hihara S, Pack CG, Kaizu K, Tani T, Hanafusa T, Nozaki T, et al. Local nucleosome dynamics facilitate chromatin accessibility in living mammalian cells. Cell Rep. 2012;2(6):1645-56. doi: 10.1016/j.celrep.2012.11.008. PubMed PMID: 23246002.

22. Nozaki T, Kaizu K, Pack CG, Tamura S, Tani T, Hihara S, et al. Flexible and dynamic nucleosome fiber in living mammalian cells. Nucleus. 2013;4(5):349-56. doi: 10.4161/nucl.26053. PubMed PMID: 23945462; PubMed Central PMCID: PMCPMC3899123.

23. Maeshima K, Ide S, Hibino K, Sasai M. Liquid-like behavior of chromatin. Curr Opin Genet Dev. 2016;37:36-45. doi: 10.1016/j.gde.2015.11.006. PubMed PMID: 26826680.

24. Shinkai S, Nozaki T, Maeshima K, Togashi Y. Dynamic Nucleosome Movement Provides Structural Information of Topological Chromatin Domains in Living Human Cells. PLoS Comput Biol. 2016;12(10):e1005136. doi: 10.1371/journal.pcbi.1005136. PubMed PMID: 27764097; PubMed Central PMCID: PMCPMC5072619.

25. Lakowicz JR. Principles of Fluorescence Spectroscopy. 3 ed: Springer US; 2006.

26. Kubo R, Toda M, Hashitsume N. Statistical Physics II: Nonequilibrium Statistical Mechanics. 2 ed: Springer-Verlag Berlin Heidelberg; 1991.

27. Nakane Y, Sasaki A, Kinjo M, Jin T. Bovine serum albumin-coated quantum dots as a cytoplasmic viscosity probe in a single living cell. Analytical Methods. 2012;4(7):1903. doi: 10.1039/c2ay25318f.

28. Doi M, Edwards SF. The Theory of Polymer Dynamics: Oxford University Press; 1988.

29. Tokunaga M, Imamoto N, Sakata-Sogawa K. Highly inclined thin illumination enables clear single-molecule imaging in cells. Nat Meth. 2008;5(2):159-61. doi: 10.1038/nmeth1171.

30. Subach FV, Patterson GH, Manley S, Gillette JM, Lippincott-Schwartz J, Verkhusha VV. Photoactivatable mCherry for high-resolution two-color fluorescence microscopy. Nat Meth. 2009;6(2):153-9. doi: 10.1038/nmeth.1298.

31. Croft JA, Bridger JM, Boyle S, Perry P, Teague P, Bickmore WA. Differences in the Localization and Morphology of Chromosomes in the Human Nucleus. The Journal of Cell Biology. 1999;145:1119-31.

32. Solovei I, Kreysing M, Lanctot C, Kosem S, Peichl L, Cremer T, et al. Nuclear architecture of rod photoreceptor cells adapts to vision in mammalian evolution. Cell. 2009;137(2):356-68. doi: 10.1016/j.cell.2009.01.052. PubMed PMID: 19379699.

33. Bickmore WA, van Steensel B. Genome architecture: domain organization of interphase chromosomes. Cell. 2013;152(6):1270-84. doi: 10.1016/j.cell.2013.02.001. PubMed PMID: 23498936.

